# Succinylation of a KEAP1 sensor lysine promotes NRF2 activation

**DOI:** 10.1101/2023.05.08.539908

**Authors:** Lara Ibrahim, Caroline Stanton, Kayla Nutsch, Thu Nguyen, Chloris Li-Ma, Yeonjin Ko, Gabriel C. Lander, R. Luke Wiseman, Michael J. Bollong

## Abstract

Crosstalk between metabolism and stress-responsive signaling is essential to maintaining cellular homeostasis. One way this crosstalk is achieved is through the covalent modification of proteins by endogenous, reactive metabolites that regulate the activity of key stress-responsive transcription factors such as NRF2. Several metabolites including methylglyoxal, glyceraldehyde 3-phosphate, fumarate, and itaconate covalently modify sensor cysteines of the NRF2 regulatory protein KEAP1, resulting in stabilization of NRF2 and activation of its cytoprotective transcriptional program. Here, we employed a shRNA-based screen targeting the enzymes of central carbon metabolism to identify additional regulatory nodes bridging metabolic pathways to NRF2 activation. We found that succinic anhydride, increased by genetic depletion of the TCA cycle enzyme succinyl-CoA synthetase or by direct administration, results in N-succinylation of lysine 131 of KEAP1 to activate NRF2 transcriptional signaling. This study identifies KEAP1 as capable of sensing reactive metabolites not only by several cysteine residues but also by a conserved lysine residue, indicating its potential to sense an expanded repertoire of reactive metabolic messengers.

## Introduction

The accumulation of intracellular reactive oxygen species (ROS), also referred to as oxidative stress, leads to the modification of cellular macromolecules including proteins, lipids, and nucleic acids and induces cellular dysfunction implicated in the onset and pathogenesis of many diseases.^1^ Crucial to counteracting the damaging effects of ROS is the inducible transcriptional activity of nuclear factor erythroid 2-related factor 2 (NRF2).^2^ This cap’n’collar (CNC) basic leucine zipper transcription factor (TF) serves as the primary stress-responsive transcription factor activated in response to oxidative stress and is responsible for inducing expression of antioxidant gene products to mitigate ROS-associated damage.^3^ In unstressed cellular conditions, NRF2 is sequestered in the cytoplasm and constitutively ubiquitinated through interactions with its repressor protein and Cullin 3 (CUL3) E3 ligase adaptor KEAP1 (Kelch-like ECH-associated protein).^4^ In the presence of electrophilic xenobiotics or increased ROS, covalent modification of “sensor” cysteines in KEAP1 result in reduced NRF2 ubiquitination and degradation.^5^ This stabilization allows NRF2 to translocate into the nucleus where it binds to antioxidant response elements (AREs) in the regulatory sequences of target genes involved in the maintenance of cellular redox, such as NAD(P)H quinone dehydrogenase 1 (NQO1). Dysregulated KEAP1-NRF2 activity is implicated in the pathogenesis of a wide range of diseases including neurodegeneration^5^, cancer^6^, inflammation^7^, and diabetes^8^. Accordingly, understanding the processes regulating KEAP1-NRF2 activity is critical for developing therapeutics strategies for treating a host of age-related diseases.

The modification of the NRF2-repressor and electrophile-sensor KEAP1 is of particular interest, as recent results have highlighted unique links between metabolism and KEAP1 modification in response to cellular stress. Numerous endogenous metabolites have been identified as signaling molecules, acting to directly modify KEAP1 ‘sensor’ cysteines via covalent non-enzymatically derived posttranslational modifications. A common target residue is cysteine 151 within the BTB domain of KEAP1, which serves as a scaffold for the assembly of the CUL3-based E3 ligase complex that targets NRF2 for degradation.^9^ Fumarate, an oncometabolite from the TCA cycle, has been shown to modify cysteine 151 of KEAP1 via S-succinylation-based modification.^10, 11^ Renal cell carcinomas often have mutations in fumarate hydratase, leading to accumulation of fumarate, resulting in enhanced NRF2 activity and tumor progression.^10^ Another TCA-derived metabolite, itaconate, has been found to alkylate cysteine 151 in KEAP1 and activate NRF2.^12, 13^ Intriguingly, the anti-inflammatory capacity of itaconate in macrophages requires the obligate activation of NRF2, highlighting the functional importance of itaconate-dependent KEAP1 modification.

More recently, evidence of reactive metabolites serving as signaling molecules to the KEAP1-NRF2 axis has come from two unbiased, high throughput reporter-based screens which identified non-covalent pharmacological activators of NRF2. The identified small molecules, CBR-470-1^14^ and sAKZ692^15^, were found to inhibit the glycolytic enzymes phosphoglycerate kinase 1 (PGK1) and pyruvate kinase (PKM2) resulting in the accumulation of the glycolytic metabolites methylglyoxal (MGO) and glyceraldehyde 3-phosphate (Ga3P), respectively. Mechanistic analyses indicated that these metabolites activate NRF2 through distinct mechanisms, involving two previously unidentified, non-enzymatically derived post translational modifications of KEAP1. MGO accumulation results in intramolecular crosslinks between cysteine 151 and arginine 15 or arginine 135 of neighboring KEAP1 molecules, inducing covalent KEAP1 dimerization and increased NRF2 activity.^14^ In contrast, Ga3P induces S-lactoylation of cysteine 273 of KEAP1, which stabilizes NRF2 and promotes its transcriptional activation.^15^ These results highlight key links between metabolic pathways and KEAP1-NRF2 signaling that are important for broadly coordinating metabolic and antioxidant regulation in response to diverse types of cellular insults.

The above results indicate that due to the virtue of their chemical structures or their positions within a given metabolic pathway, some reactive metabolites have most likely been evolutionarily selected to relay metabolic information to the stress-sensing machinery of the cell. Given the observed capacity of several central metabolites to modify KEAP1 covalently, we hypothesized that several other conserved nodes throughout central carbon metabolism might also signal to the oxidative stress sensing machinery. To test this hypothesis, we performed an shRNA screen of genes of the TCA cycle, glycolysis, the pentose phosphate pathway, and the glyoxalase system to identify if decreasing a specific enzymatic activity of one of these enzymes might increase the levels of a KEAP1 reactive metabolite capable of activating NRF2. Here, we show that genetic perturbations of the TCA cycle promote NRF2 activation through the non-enzymatic N-succinylation of a regulatory lysine in the BTB domain of KEAP1. These results demonstrate a new posttranslational mechanism by which KEAP1 can sense metabolic disruptions, further linking the status of cellular metabolic health to the activation of the NRF2 antioxidant signaling pathway.

## Results

### Succinyl CoA synthetase depletion activates NRF2 transcriptional signaling

To identify potential metabolic regulators of NRF2 activity, we screened shRNAs against known metabolic enzymes to identify those whose depletion induced NRF2 activation. Towards that aim, we transduced K562 lymphoblast cells with the NRF2 reporter plasmid ARE-GFP-LUC, which encodes the NRF2 ARE binding site from the promoter of the human *NQO1* gene driving the expression of GFP and luciferase (**Fig. 1A**). This enables measurement of NRF2 activation by GFP fluorescence or by luciferase activity. Monoclonal cells expressing this reporter were then transduced with shRNA targeting *KEAP1*, and after 72 hrs, we selected the cell line with the most dynamic and reproducible induction of GFP signal measured by flow cytometry. We then transduced reporter cells with shRNAs targeting genes encoding 30 metabolic enzymes (3 shRNAs/gene) of glycolysis, the pentose phosphate pathway (PPP), the glyoxalase system, or the TCA cycle (**Fig. 1B**). We then measured ARE-GFP reporter activity by flow cytometry (**Fig. 1C**). This approach identified genes whose depletion are known to increase intracellular levels of reactive metabolites capable of activating NRF2. For example, depletion of fumarate hydratase (FH) was identified as a top hit in the screen, activating the ARE-GFP reporter, consistent with studies showing fumarate hydratase deficiency increases cellular levels of the KEAP1-reactive metabolite fumarate.^10, 11, 16^ Further, depletion of transketolase (TKT) or transaldolase (TALDO1), two enzymes in the non-oxidative branch of the PPP, also resulted in ARE-GFP reporter activation. This result likely reflects the ability for these enzymes to reversibly catalyze reactions with the glycolytic metabolite Ga3P, which we have previously shown to activate NRF2 through direct modification of KEAP1.^15^ Collectively, these results confirm the ability of our approach to identify links between perturbations of metabolic pathways and NRF2 activation.

**Figure 1.**
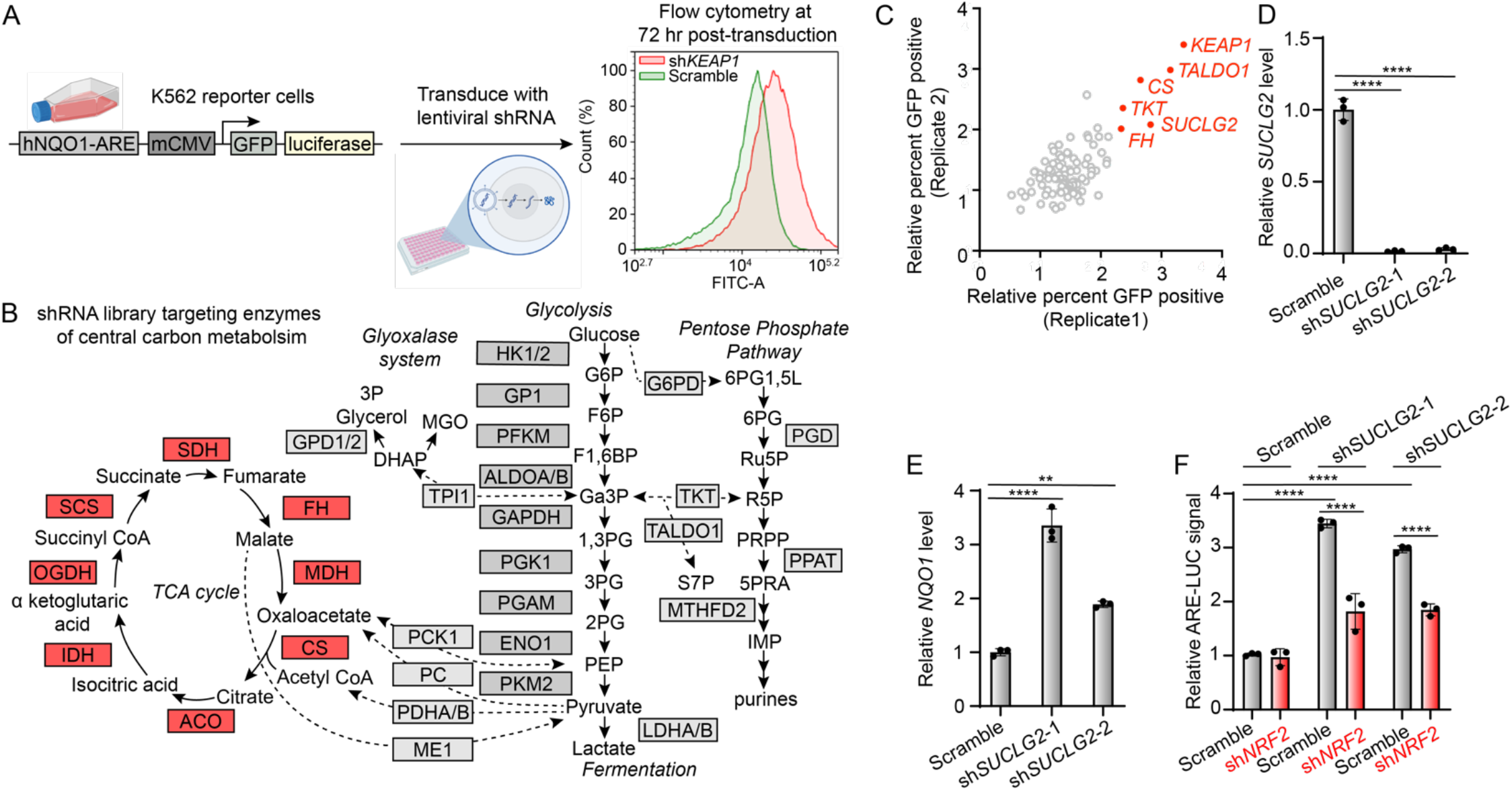
Loss of succinyl coA synthetase activates NRF2 driven transcription. A) Schematic of the cellular screening platform using ARE-GFP-LUC K562 reporter cells. Shown is a representative plot indicating positivity for GFP after transduction with an shRNA targeting *KEAP1*. B) Schematic depicting the enzymes of central carbon metabolism targeted by the shRNA screen (boxes). C) Replicate fold increases in percent GFP positive ARE-GFP-LUC K562 cells after a 72 hr exposure to the indicated shRNAs of the screen. D,E) Relative transcript levels of *SUCLG2* (D) and *NQO1* (E) after 48 hr transient transfection of HEK293T cells with shRNAs targeting *SUCLG2* (*n*=3, ****P<0.0001, **P<0.01, one-way ANOVA). F) Relative luminescence values of ARE-LUC reporter activity 72 hr after transduction of lentiviruses encoding shRNAs to *SUCLG2* and *NRF2* in ARE-GFP-LUC K562 (*n*=3, ****P<0.0001, two-way ANOVA).

Interestingly, depletion of *SUCLG2*, a subunit of succinyl coA synthetase (SCS) enzyme complex of the mitochondrial TCA cycle also activated the ARE-GFP reporter (**Fig. 1C**). The magnitude of reporter activation, like that induced by depletion of other enzymes, was dependent on the amount of transduced lentiviral shRNA, indicating dose-dependent activation (**Fig. S1A**). Depletion of *SUCLG2* by two distinct shRNAs was confirmed by qPCR in HEK293T cells (**Fig. 1D**). We demonstrated that depletion of *SUCLG2* increased expression of the NRF2 target gene *NQO1* in HEK293T cells by qPCR (**Fig. 1E**). Further, *SUCLG2* depletion was also shown to increase luciferase activity in K562 cells expressing this dual reporter (**Fig. 1A, F**). Knockdown of *NRF2* inhibited the increase in ARE-luciferase activity induced by *SUCLG2* depletion, confirming that this activation reflects NRF2-depedent transcriptional activation (**Fig. 1F**). The ability of *SUCLG2* depletion to activate reporter activity was observed in cells grown in glucose or galactose containing media (**Fig. S1B**), indicating that this effect is independent of the source of cellular ATP. Collectively, these results show that depletion of *SUCLG2* activates NRF2 transcriptional signaling, revealing a new link between mitochondrial TCA activity and NRF2 activation.

### Succinic anhydride activates NRF2

We next sought to identify if increased levels of a reactive metabolite were responsible for NRF2 activation induced by *SUCLG2* depletion. We demonstrated that ARE-luciferase activity induced by *SUCLG2* depletion is decreased by treatment of K562 reporter cells with the thiol-containing antioxidant glutathione (GSH; **Fig. 2A**). However, the general antioxidant vitamin E did not reduce ARE-luciferase reporter activity, while the mitochondrial-targeted antioxidant MitoTEMPO showed only a mild reduction in reporter activation. These results are consistent with a model in which *SUCLG2* depletion increases concentration of a reactive metabolite that could activate NRF2 through KEAP1 modification.

**Figure 2.**
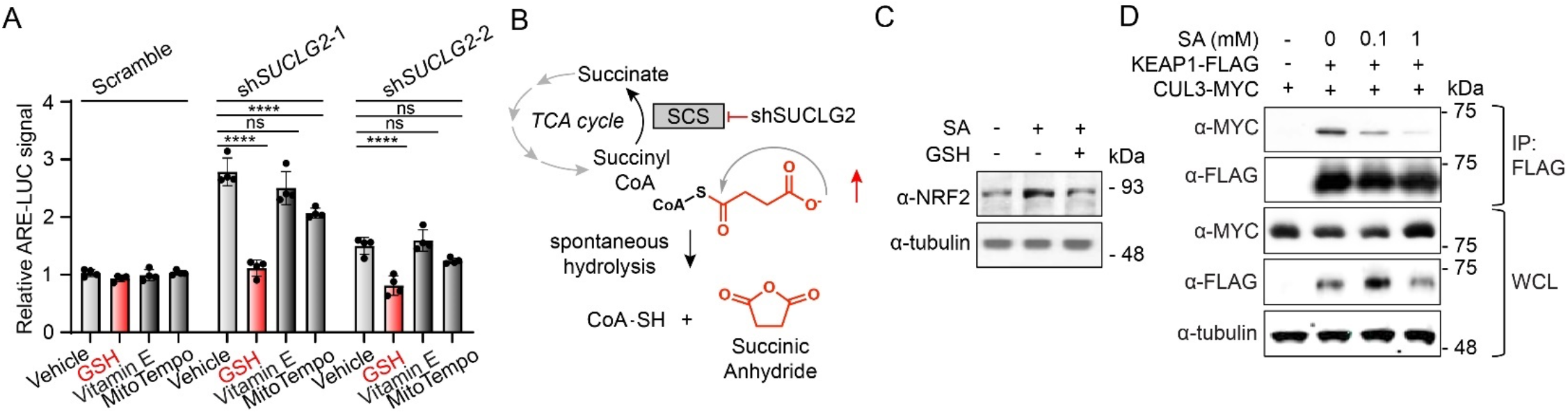
Augmented levels of succinic anhydride activate NRF2 signaling. A) Relative luminescence signal of ARE-LUC reporter activity induced shRNAs targeting *SUCLG2* in ARE-GFP-LUC K562 cells in the presence of the indicated antioxidants (10 mM each; *n*=4, ns = not significant P>0.05, ****P<0.0001, two-way ANOVA). B) Schematic depicting the increase of succinyl CoA and subsequent spontaneous formation of SA after knockdown of the SCS subunit *SUCLG2*. C) Representative Western blotting for NRF2 levels after 4 hrs treatment of HEK29T cells with SA (10 mM) in the absence or presence of GSH (10 mM). D) Representative anti-MYC Western blotting of anti-FLAG coimmunoprecipitated material after 1 hr of SA treatment in HEK293T expressing MYC-CUL3 and KEAP1-FLAG transgenes.

Depletion of SCS leads to increased levels of succinyl CoA and global protein N-succinylation.^17^ Further, reactive acyl-CoA metabolites are known to traverse the mitochondrial membranes and non-enzymatically acylate and succinylate lysines and cysteines on cytosolic proteins.^18, 19^ This suggests that increases in succinyl-CoA induced by *SUCLG2* depletion could contribute to NRF2 activation observed under these conditions (**Fig. 2B**). Previous studies indicate that succinyl-CoA self-hydrolyzes to generate free coenzyme A and the highly reactive cyclic byproduct succinic anhydride, which has been shown to be the most likely mechanism of protein succinylation.^20^ We hypothesized that knockdown of *SUCLG2* leads to increased levels of succinic anhydride (SA) that modify KEAP1 and subsequently activates NRF2 signaling (**Fig. 2B**). Interestingly, treatment with SA stabilized NRF2 in HEK293T cells, while co-treatment with GSH decreased this stabilization (**Fig. 2C**). To further define the impact of SA on NRF2 stabilization, we monitored the association between KEAP1-FLAG and MYC-CUL3 – the ligase that targets NRF2 for degradation – in HEK293T cells. We observed dose-dependent reductions in recovered MYC-CUL3 in KEAP1-FLAG immunoprecipitations, indicating that SA disrupted the interaction between KEAP1 and CUL3 (**Fig. 2D**). Treatment with SA also activated the ARE reporter in K562 cells (**Fig. S2A, B**). Importantly, neither *SUCLG2* knockdown nor SA treatment increased reactive oxygen species (ROS) in K562 cells, as measured by DCFDA (**Fig. S2C**). These results indicate that SA activates NRF2 signaling through a mechanism independent of inducing oxidative stress, instead likely acting by covalently modifying KEAP1.

### SA increases lysine succinylation on KEAP1

We next sought to identify the potential post translational modification of KEAP1 formed by SA. We employed a succinic anhydride probe molecule containing an alkyne reactivity handle (SA-alkyne) compatible with click chemistry-based conjugation to monitor KEAP1 modification (**Fig. 3A**). We confirmed that SA-alkyne activates the ARE-luciferase reporter in IMR32 cells to a similar extent to that observed with SA (**Fig. 3B**). Treatment of HEK293T cells overexpressing KEAP1-FLAG with increasing concentrations of SA-alkyne, significantly increased the accumulation of modified KEAP1 identified in anti-FLAG based immunoprecipitation experiments (**Fig. 3C**). Critically, SA-alkyne modification of KEAP1 was competed by increasing concentrations of SA, confirming that SA covalently modifies KEAP1 (**Fig. 3D**).

**Figure 3.**
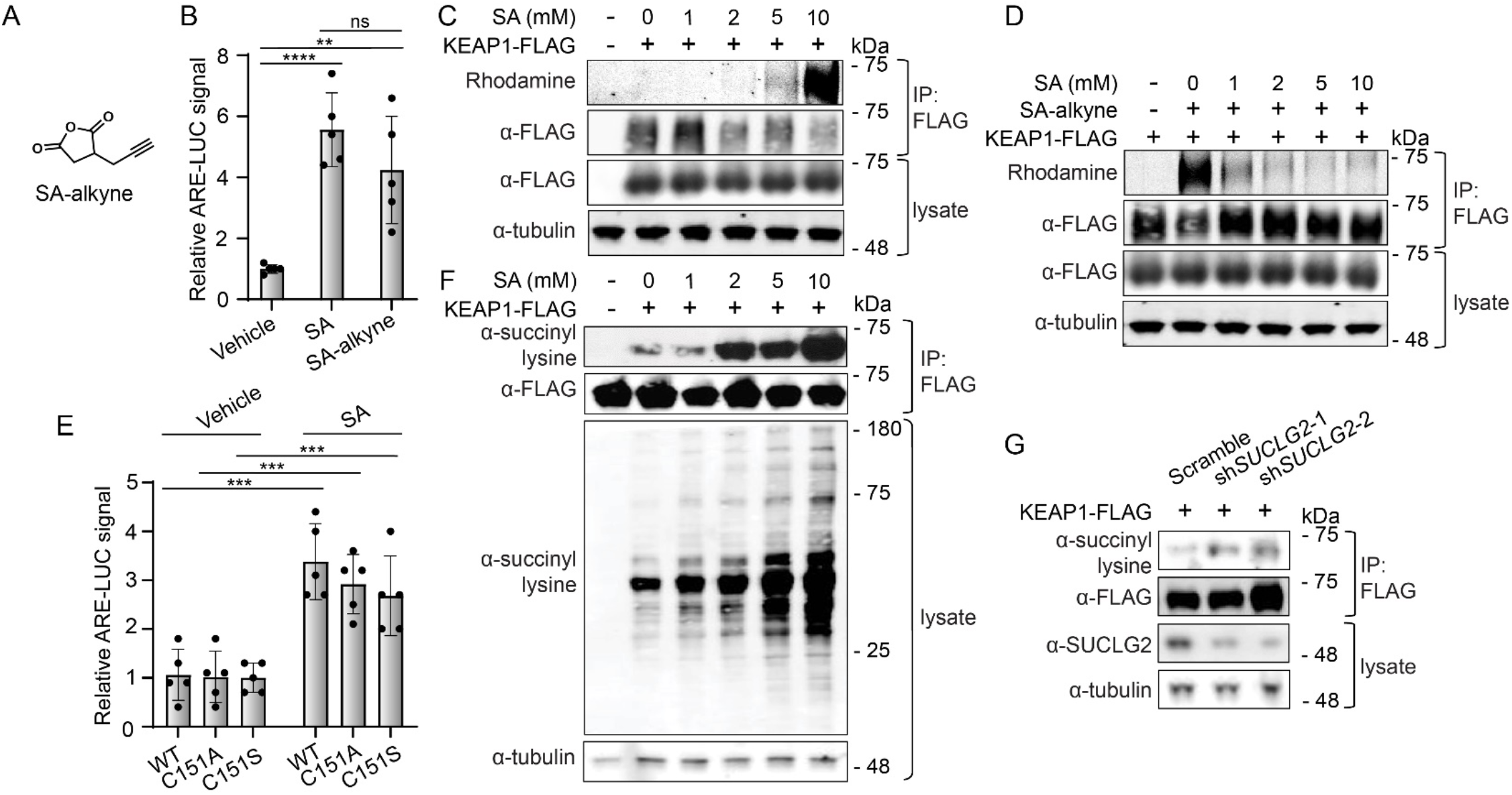
SA accumulation leads to covalent modification of KEAP1 lysines. A) Structure of succinic anhydride alkyne (SA-alkyne). B) Relative luminescence values of ARE-LUC reporter activity from IMR32 cells treated with SA or SA-alkyne (12 mM) for 4 hrs (*n*=5, ****P<0.0001, **P<0.01, one-way ANOVA). C) Representative fluorescence scan of rhodamine azide labeling of anti-FLAG immunoprecipitated material after treatment of HEK293T cells expressing KEAP1-FLAG with the indicated concentrations of SA-alkyne (10 mM) for 1 hr. D) Representative fluorescence scan of rhodamine azide labeling of anti-FLAG immunoprecipitated material after pretreated with the indicated concentrations of SA followed by a 1 hr labeling with SA-alkyne (10 mM) in HEK293T cells expressing KEAP1-FLAG. E) Relative ARE-LUC activity from IMR32 cells expressing the indicated KEAP1 mutant and MYC-NRF2 and then treated with 1 mM SA for 4 hrs (*n*=5, ***P<0.001, two-way ANOVA). F) Western blotting for anti-N-succinyl lysine positivity of anti-FLAG immunoprecipitated material from HEK293T cells expressing KEAP1-FLAG after 1 hr treatment with the indicated concentrations of SA. G) Western blotting for anti-N-succinyl lysine positivity of anti-FLAG immunoprecipitated material from HEK293T cells expressing KEAP1-FLAG and shRNAs targeting *SUCLG2*, stabilized by the desuccinylase inhibitor ET-29 (10 μM).

KEAP1 activity is primarily regulated through covalent modification of cysteine residues, most notably cysteine 151. To determine whether SA activates NRF2 through covalent modification of cysteine 151, we overexpressed wild-type KEAP1-FLAG or C151A/S KEAP1-FLAG in IMR32 cells and monitored ARE-luciferase activation. Surprisingly, we found that mutation of C151 did not disrupt SA-dependent ARE-luciferase activation, suggesting that SA modifies KEAP1 on a different residue (**Fig. 3E**). SA has previously been shown to modify lysine residues in proteins. To determine if SA also modifies lysine residues within KEAP1, we employed an antibody that detects lysine N-succinylation, evaluating if anti-FLAG immunoprecipitated KEAP1-FLAG displayed immunopositivity to increasing SA levels. Not only did exogenous SA treatment increase global succinylation of proteins, as anticipated, (**Fig. 3F**), we also observed dose-dependent increases of lysine succinylation in KEAP1-FLAG immunopurified from SA-treated cells (**Fig. 3F**). *SUCLG2* depletion also increased lysine succinylation on KEAP1-FLAG in HEK293T cells co-treated with ET-29 (**Fig. 3G**) - an inhibitor of the principle intracellular lysine desuccinylase SIRT5 that stabilizes succinylated lysines (**Fig. S3A**).^21^ Further, we found that treatment of HEK293T cells with SA increased lysine succinylation of endogenous KEAP1 (**Fig. S3B**). Similar results were observed in reactions with recombinant KEAP1 and either SA or succinyl-coA (**Fig. S3C, D**).

### Lysine 131 succinylation functionally inhibits KEAP1 activity

The above results demonstrate that treatment with SA or *SUCLG2* depletion increases lysine succinylation on KEAP1. Next, we wanted to determine if this modification was functionally important to NRF2 activity. To identify the specific lysine residues of KEAP1 modified by SA, we used MS/MS analysis of KEAP1-FLAG immunoprecipitated from HEK293T cells treated with SA (**Fig. 4A, Fig. S4**). This effort identified lysine 131 of the BTB domain of KEAP1 as the principal lysine modified by this metabolite. Lysine succinylation of residue 131 was also observed from recombinant KEAP1 preparations treated with SA (**Fig. S5A, B**). Importantly, overexpression of K131A KEAP1 decreased labeling by SA-alkyne (**Fig. 4B**) and reduced KEAP1 lysine succinylation in SA-treated HEK293T cells (**Fig. 4C**). Overexpressed KEAP1 containing mutations in the key electrophile sensing residue C151 did not affect these modifications. This indicates that K131 is the primary site of lysine succinylation on KEAP1.

**Figure 4.**
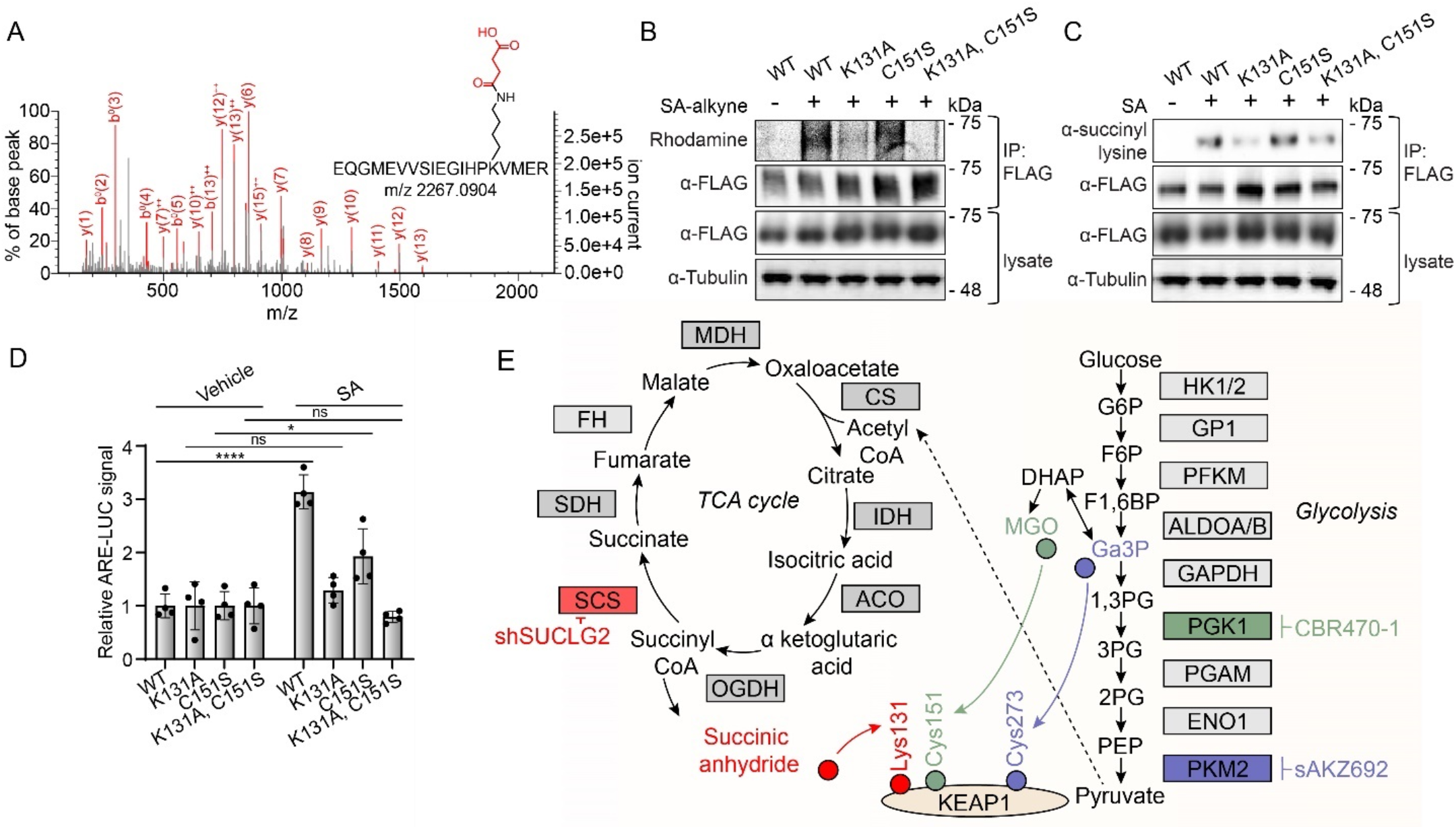
N-succinylation of Lysine 131 functionally inhibits KEAP1 activity. A) MS/MS spectra of the N-succinylated K131 containing tryptic peptide immunoprecipitated from overexpressed KEAP1-FLAG from HEK293T cells treated with SA (10 mM). B) Representative fluorescence scan of rhodamine azide labeling of anti-FLAG immunoprecipitated material after pretreated with the indicated concentrations of SA followed by a 1 hr labeling with SA-alkyne (5 mM) in HEK293T cells expressing the indicated KEAP1-FLAG transgenes. C) Western blotting for anti-N-succinyl lysine positivity of anti-FLAG immunoprecipitated material from HEK293T cells expressing the indicated KEAP1-FLAG transgene mutants after 1 hr treatment with SA (1 mM). D) Relative ARE-luciferase reporter activity from IMR32 cells expressing the indicated KEAP1-FLAG mutant transgenes and then treated with 1 mM SA for 4 hrs (*n*=4, ns = not significant P>0.05; ****P<0.0001, *P<0.05, two-way ANOVA). E) Schematic depicting metabolites from central carbon metabolism that can covalently modify the indicated residues on KEAP1.

Although K131 exists in a flexible disordered region that exhibits weak electron density in structural models^22^, this lysine residue appears uniquely positioned within a basic pocket that surrounds the key electrophile sensor reside C151 (**Fig. S5C**).^23^ This suggests that K131 succinylation may directly impact KEAP1 activity, and thus NRF2 activation. Consistent with this, we found that K131A KEAP1 blocked SA-dependent activation of the ARE-luciferase reporter, while C151S KEAP1 did not (**Fig. 4D**). This contrasts with established NRF2 activators like bardoxolone that activate NRF2 through C151 modification (**Fig. S5D**). Collectively, these results indicate that SA-dependent succinylation of KEAP1 at residue K131 regulates NRF2 activation.

## Discussion

To comprehensively map the interactions between reactive metabolites and the oxidative stress sensing machinery of the cell, here we have conducted an shRNA screen of the enzymes in central carbon metabolism, targeting genes involved in glycolysis, the TCA cycle, the PPP, and the glyoxalase system. This effort identified the mitochondrial enzyme succinyl coA synthetase (SCS) as a key regulator of NRF2 activity, as genetic depletion of SCS leads to the N-succinylation of KEAP1 lysine 131 and subsequent NRF2 activation. The human KEAP1 protein contains 27 cysteines with at least 11 of them having been shown to play sensing roles, capable of being alkylated by exogenous electrophilic chemicals, resulting in the decreased ubiquitination of NRF2.^9^ The molecular details by which covalent modification results in NRF2 activation have remained elusive; however, the consensus from the field suggests that relatively small posttranslational modifications of these cysteines most likely results in profound conformational changes in the KEAP1 intermolecular complex.^24^ Here, we add evidence suggesting that in addition to this canonical cysteine sensing paradigm, that modifications to other sensor residues, like the lysine modification identified here, can additionally result in functional NRF2 activation.

Among the most targeted sensor cysteines in KEAP1 is cysteine 151 of the BTB domain, which has been shown to be covalently modified by numerous synthetic small molecules, including the clinically investigated oleananes Bardoxolone and Omaveloxelone.^25^ Notably, lysine 131 occupies a proximal site to cysteine 151 across the Bardoxolone binding cleft (6.3 Å apart in the crystal structure 4CXI).^22^ Lysine 131, along with neighboring residues arginine 135 and lysine 150 have been shown to act as bases, locally repressing the pKa of cysteine 151, enabling its potent sensing of electrophilic chemicals.^23^ To the extent that these neighboring basic residues may also increase the nucleophilicity of lysine 131 and enable its ability to sense electrophiles will necessarily be the work of future crystallographic and biochemical studies.

We have shown that lysine 131 modification in response to *SUCLG2* depletion is derived from the reactive byproduct of succinyl-CoA degradation the metabolite, succinic anhydride (SA). Importantly, several other mitochondrial metabolites have been shown to modify KEAP1, including fumarate and itacontate, both of which, like SA, are derived from the TCA cycle. Interestingly, KEAP1 has been shown to exist in two sensing forms, a cytoplasmic complex and one that is tethered to the outer face of the mitochondrial membrane via interactions with PGAM Family Member 5 (PGAM5). ^26^ These observations raise the intriguing possibility that this ternary complex of KEAP1, NRF2, and PGAM5 may provide a molecular sensing outpost for relaying specific mitochondrial metabolic information to the oxidative stress response.

The TCA cycle is a tightly regulated metabolic pathway, using several feedback loops to maintain cellular homeostasis. Interestingly, succinyl-CoA levels have been shown to regulate metabolic flux of the TCA cycle by inhibiting citrate synthase and α-ketoglutarate dehydrogenase through a negative feedback mechanism.^27^ To the extent that increased succinyl-CoA levels might additionally serve as a sentinel for signaling TCA cycle imbalance to the oxidative stress sensing machinery will be the work of future studies. In addition to SCS, we identified another TCA cycle enzyme as a hit from our screen, as genetic depletion of citrate synthetase (CS) was found to result in mean increase of 2.7-fold in ARE reporter activity (**Fig. 1C**). CS catalyzes the formation of citrate from oxaloacetate (OA), a high energy carbonyl containing metabolite that is the product of the TCA cycle.

Whether OA, and perhaps other carbonyl containing metabolites of the TCA cycle might additionally be sensed by KEAP1 via posttranslational modifications of cysteines or lysines will be of keen interest in future work. Projecting forward, this work will provide the basis for understanding how these and other molecules maintain cellular homeostasis. Likewise, the novel lysine sensing mechanism described here might further inform efforts to covalently target KEAP1 with greater selectivity, potentially broadening the therapeutic scope of therapeutics aimed at activating NRF2 with increased tolerability.

## Acknowledgements

We thank Linh Truc Hoang of the Scripps Research Center for Metabolomics for assistance with mass spectrometry related experiments, and Zoe Adams of the Dawson Lab at Scripps Research for help with peptide synthesis. We also thank Nelson Wu of the Schief Lab at Scripps Research for advice regarding structural modeling. This work was supported by the NIH (GM146865 to MJB, DK107604 and AG046495 to RLW).

## Author contributions

LI, RLW, and MJB designed research. LI, TN, and CLM performed biochemical and cell-based biological experiments. KN and YK purified recombinant protein. CS conducted mass spectrometry. LI analyzed data. LI, RLW, and MJB wrote the paper. Financial support by the Skaggs Institute for Chemical Biology at Scripps Research.

## Figures

**Supplementary Figure 1.**
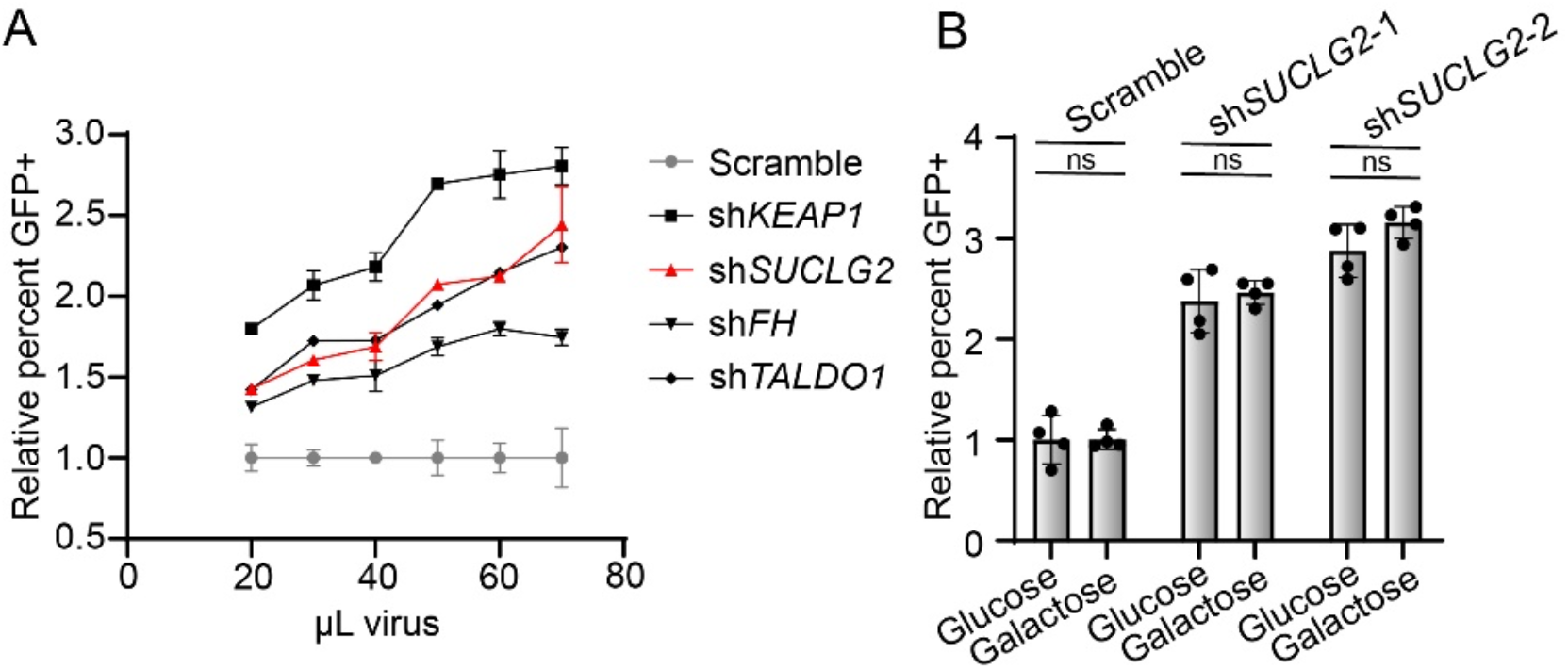
Knockdown of *SUCLG2* activates ARE-GFP-LUC. A) Relative ARE activation from ARE-GFP-LUC K562 cells in response to the indicated volumes of lentiviruses encoding the indicated shRNAs. B) Relative ARE activation from ARE-GFP-LUC K562 cells in response to treatment with lentiviruses encoding the indicated shRNAs to *SUCLG2* grown in either glucose or galactose (*n=*4; ns = not significant *P*>0.05, two-way ANOVA).

**Supplementary Figure 2.**
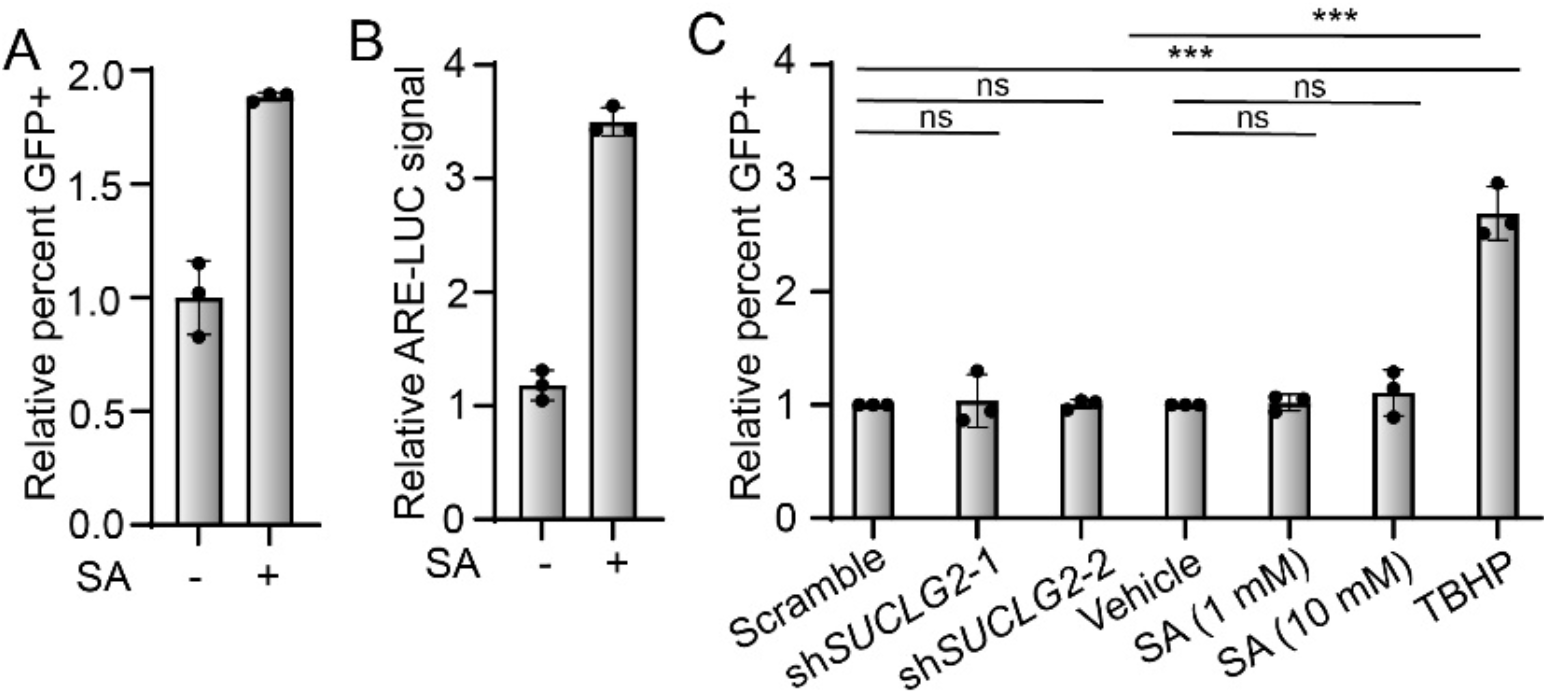
SA activates ARE-GFP-LUC in K562 cells. A) Relative percent GFP positive K562 ARE-GFP-LUC cells after treatment with 50 mM SA for 4 hrs (*n*=3, ***P<0.001, t-test). B) Relative luminescence signal of reporter activity from K562 ARE-GFP-LUC cells after treatment with 50 mM SA for 4 hrs (*n*=3, ***P<0.001, t-test). C) Relative percent GFP positive measurements indicating the presence of reactive species from K562 cells treated with virus expressing shRNAs targeting *SUCLG2* or treatment with the indicated compounds (TBHP = tert-butyl hydro peroxide; *n*=3, ns = not significant P>0.05, ****P<0.0001, one-way ANOVA).

**Supplementary Figure 3.**
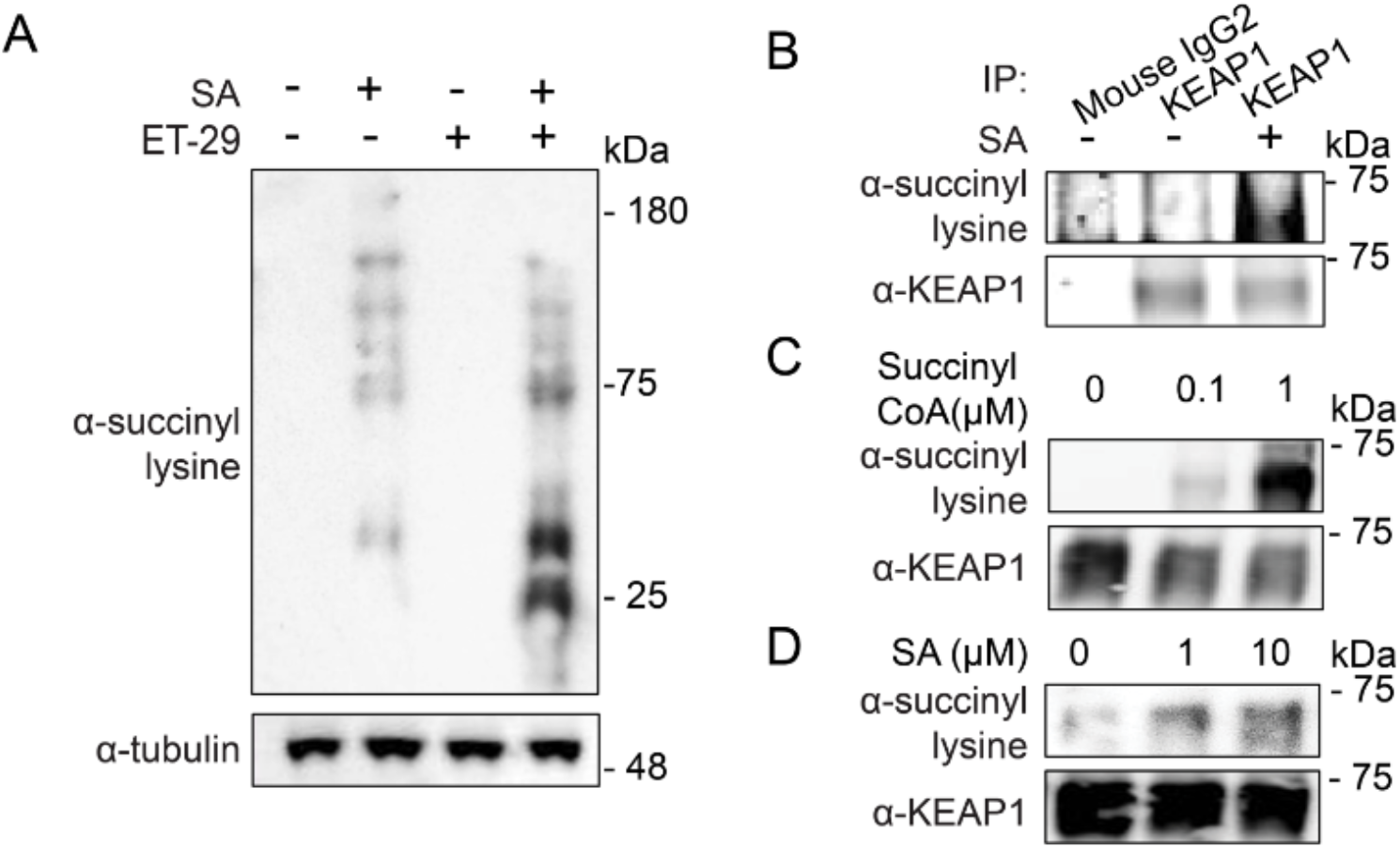
Covalent modification of KEAP1 lysines in response to SA. A) Western blotting analysis for anti-succinyl lysine positivity after treatment of HEK293T cells with ET-29 (10 μM) and SA (5 mM, 1 hr). B) Western blotting anti-succinyl lysine positivity from pulldowns of endogenous KEAP1 from HEK293T cells treated with SA (5 mM, 1 hr). Western blotting anti-succinyl lysine positivity of recombinant KEAP1 after a 1 hr in vitro treatment with the indicated concentrations of Succinyl-CoA (C) or SA (D).

**Supplementary Figure 4.**
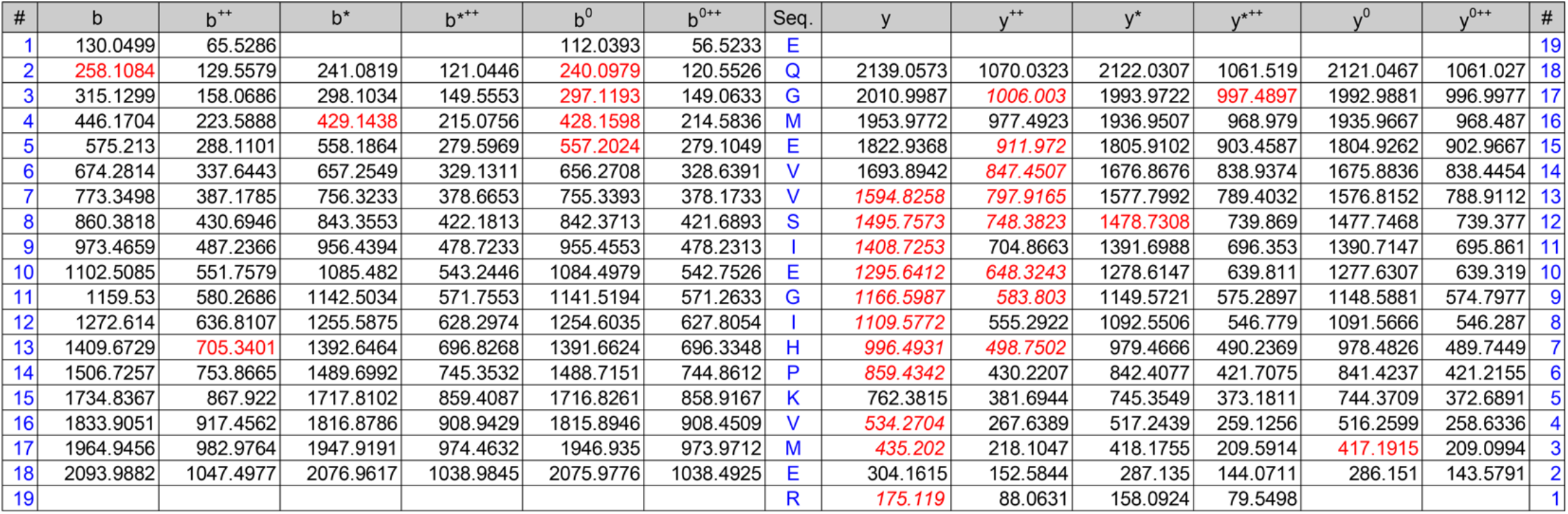
Lysine 131 is succinylated by SA in cells. *b* and *y* ion designations corresponding to MS/MS spectra in Figure 4A.

**Supplementary Figure 5:**
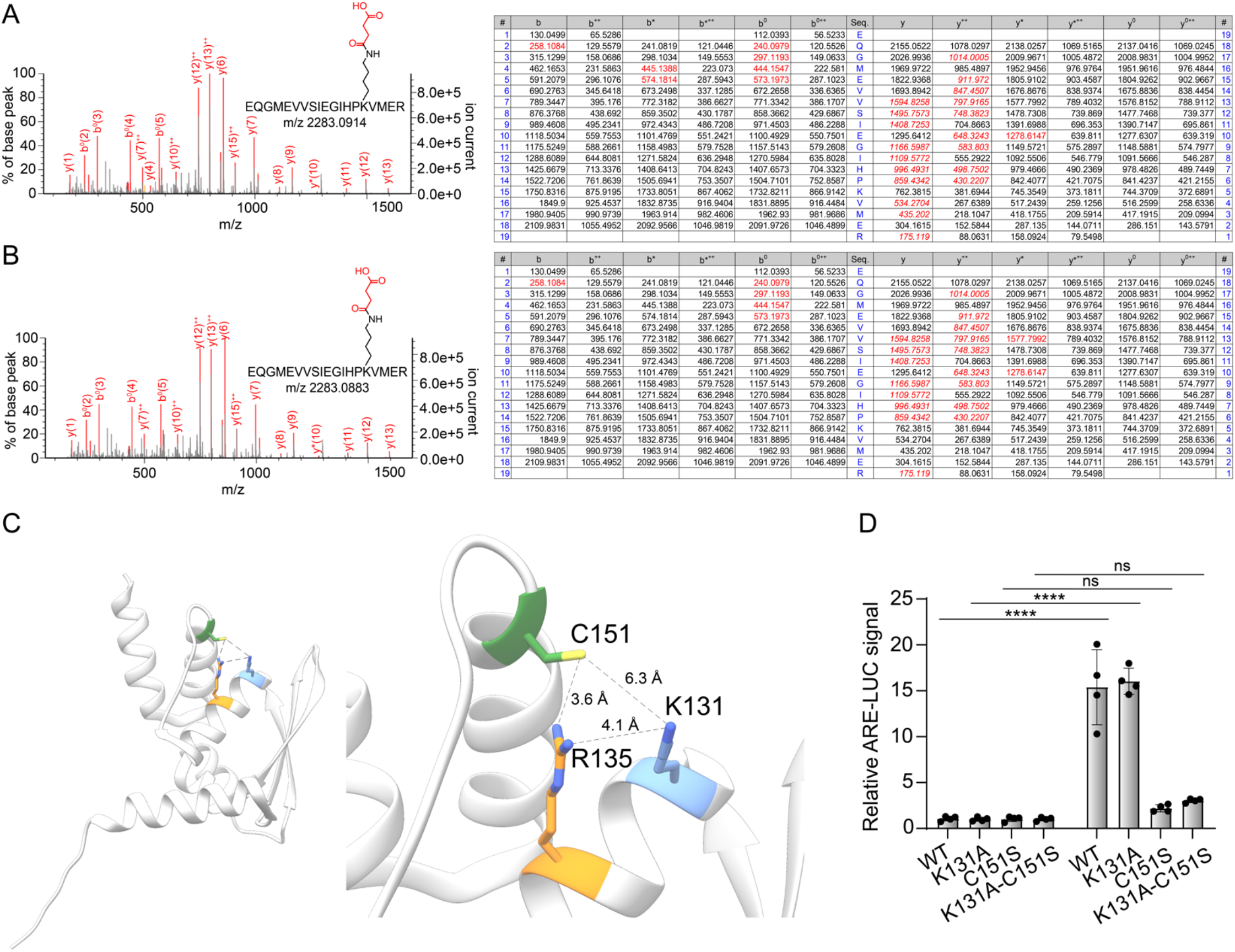
Lysine 131 of KEAP1 is covalently modified by SA. MS/MS spectra (left) along with *b* and *y* ion designations (right) corresponding to +101 Da modified tryptic KEAP1 peptides K131 from 1 hr *in vitro* reactions of SA (10 mM (A) and 1 mM (B)) with recombinant KEAP1. C) Representation of the BTB domain of KEAP1 (left; PDB: 4CXI) with inset denoting orientations of key residues K131, R135, and C151(right). D) ARE-luciferase reporter activity of IMR32 cells expressing the indicated KEAP1 transgene mutants after treatment with 500 nM bardoxolone methyl for 24 hrs (*n*=4, ns = not significant P>0.05, ****P<0.0001, two-way ANOVA).

## Methods

### Chemicals

Succinic anhydride (cat. no. 239690), TBHP (cat. no. 458139), GSH (cat. no. PHR1359), Vitamin E (cat. no. 5.00878), DCFDA (cat. no. 21884) and MitoTempo (cat. no. SML0737) were purchased from Sigma Aldrich. The succinic anhydride probe was from Enamine (cat. no. EN200-172303). ET-29 was from MedChemExpress (HY-145651).

### Cell culture

K562, HEK293T, and IMR32 cells were purchased from American Type Culture Collection (ATCC). K562 cells were maintained in RPMI (Corning) supplemented 10% with fetal bovine serum (FBS, Gibco) and 1% penicillin-Streptomycin (Pen Strep, Gibco). HEK293T and IMR32 cells were maintained in DMEM (Corning) supplemented 10% with fetal bovine serum (FBS, Gibco) and 1% penicillin-Streptomycin (Pen Strep, Gibco). For experiment with galactose media, cells were grown in DMEM (no glucose) supplemented with 10 mM galactose (Sigma Aldrich) for 1 week.

### Cloning

To generate the ARE-GFP-LUC vector, the ARE sequence from the human NQO1 promoter (CTCAGCCTTCCAAATCGCAGTCACAGTGACTCAGCAGAAT), synthesized as double stranded synthetic DNA oligo (IDT), was inserted between the EcoR1 and BglII sites in pGreenFire (System Biosciences). To generate KEAP1-FLAG mutants, single and double point mutations (C151S/A, K131A) were introduced into FLAG-KEAP1 (Addgene #28023) using primers synthesized by IDT and a Q5® Site-Directed Mutagenesis Kit (NEB).

### Lentivirus production and transduction

Lentiviruses were generated in HEK293T cells by transient expression of the indicated shRNA vectors below with psPAX2 and pMD2.G packaging vectors (Addgene plasmids 11260 and 12259). For lentiviral production in 96 well plates, 1.5*10^4^ HEK293T cells were plated in poly-d-lysine (Thermo Fisher Scientific) coated wells in 100 μL of growth medium. Transfection mixtures contained 30 μL of Optimem medium (Gibco), 100 ng of shRNA, psPAX2, and pMD2.G each, and 1μg total DNA:4μL of Fugene (Promega) per well. Viral supernatants were collected after 48 h of expression and added to target cells with the addition of a final concentration of 5 μg/mL polybrene (Sigma Aldrich).

### Stable cell line production

48 hrs after transduction of K562 cells with lentiviruses encoding the ARE-GFP-LUC construct, cultures were exposed to 5 μg/mL of puromycin for one week. Single cells were then plated in 96 well plates and allowed to expand for several weeks. The monoclonal line with the most dynamic and reproducible increase in FITC+ signal by flow cytometry after shKEAP1 transduction (tested at 48, 72, and 96 hrs post-transduction) was selected for use in the screen.

### shRNA screen in ARE-GFP-LUC K562 reporter cells

ARE-GFP-LUC K562 reporter cells (2*10^4^ in 50 μL) were dispensed into 96 well plates and transduced as described above with 100 μL of lentiviral shRNAs. After 72 hrs cells were analyzed for GFP positivity using a NovoCyte 3000 with NovoSampler Pro. Select shRNAs used to follow up hits from the original screen are shown below.

| shRNA target | Lentiviral clone from Sigma Mission shRNA |
| --- | --- |
| Non-targeting control vector | SHC002 |
| KEAP1 | TRCN0000154656 |
| FH | TRCN0000052463 |
| TALDO1 | TRCN0000052519 |
| TKT | TRCN0000054073 |
| CS | TRCN0000045703 |
| SUCLG2-1 | TRCN0000048503 |
| SUCLG2-2 | TRCN0000048505 |
| NRF2 | TRCN0000007555 |

### ARE-LUC reporter assays

IMR32 cells were plated in 40 μL of growth medium (5000 cells/well) in white 384-well plates (Greiner). The next day each well was transfected with 100 ng pTI-ARE-LUC using FuGENE HD (4 μL per ug DNA) in freshly filtered 10 μL OptiMEM medium (Gibco), or 50 ng of the reporter and 50 ng of shRNA. For experiments with KEAP1-FLAG mutants, 20 ng of MYC-NRF2 was included in the transfection mix. 24 hrs later compounds (antioxidants or SA/SA-alkyne) were added in 10 μL of 6x solutions. After treatment (24 hrs for antioxidants, 4 hrs for SA/SA-alkyne), 30 μL BrightGlo (diluted 1:3 in water) were dispensed to each well, plates shaken for one minute, and the luminescence signal recorded using an Envision plate reader (PerkinElmer).

### qRT-PCR

Treated HEK293T cells were collected from 6 well plates via trypsinization. RNA was isolated using RNeasy kits (Qiagen) and concentrations quantified using a Nanodrop instrument. 2 μg of RNA was then subjected to oligo dT-primed reverse transcription reactions (SuperScript III First Strand Synthesis Kit). Quantitative RT-PCR reactions were measured on a Viia7 Instrument (Thermo) using Clontech SYBR green-based master mix (TB Green Premix). Calculations were normalized using TUBG1 (Tubulin) as a reference. Ct values were determined using the Viia7 software and transcript abundance was calculated using the standard comparative Ct method.

### Quantification of ROS by DCFDA Fluorescence

K562 cells were plated in 96-well plates and treated/transduced as previously described. Cells were then washed and resuspended in DPBS. CM-H2DCFDA (Invitrogen) was freshly dissolved in DMSO. Cells were incubated in 5 μM CM-H2DCFDA for 30 min and analyzed on a NovoCyte 3000.

### KEAP1-FLAG labeling experiments in HEK293T cells

6-well tissue culture plates were coated with poly-d-lysine, and 5*10^5^ HEK293T cells were plated per well. Wells were transfected with 1 μg of the indicated KEAP1 encoding vectors with 4 μL Fugene HD in 0.1 mL Opti-MEM (and additional 1 μg for shSUCLG2 experiments with 8 μL Fugene HD total). After a 48 hr incubation cells were treated with the indicated concentration of SA/SA-alkyne for 1 hr. For competition experiment, cells were exposed to SA for 1 hr after initial incubation with alkyne probe. For experiment with shSUCLG2, cells were treated with ET-29 24 hrs after transfection. Cells were washed with cold PBS twice, scraped in 0.25 mL RIPA buffer containing protease inhibitor (Roche), and lysed by sonication. Insoluble materials were removed by centrifugation (13,000 g for 5 minutes at room temperature). 1 mg of lysate was immunopurified with 20 μL of M2 Anti-FLAG magnetic beads (Thermo Fisher) and FLAG tagged material eluted using an excess of FLAG peptide (Sino biological) in 50 μL. Similar procedure for co-immunoprecipitation of MYC-CUL3 with KEAP1-FLAG was followed, except without sonication. Samples were mixed with loading buffer and separated by SDS-PAGE followed by immunoblotting. For MS/MS-based proteomics, the gel was stained with Coomassie and the band containing KEAP1 was excised and submitted. For experiments with SA-alkyne, the FLAG elute from the steps above was incubated with click reagent mix (1 μL of 1.25 mM rhodamine azide in DMSO, 1 μL of 50 mM CuSO4 in H2O, 1 μL of 50 mM TCEP in water, 3 μL of 1.7 mM TBTA in tBuOH:DMSO 4:1) at room temperature for 1 hr in the dark. The reaction mixture was precipitated in cold methanol and the pellet was re-dissolved in PBS containing 0.1% SDS. The resulting sample was mixed with loading buffer and separated via SDS-PAGE. Gels were incubated in 10% EtOH for 1 hr before imaging fluorescence on a ChemiDoc instrument (Bio-Rad).

### Recombinant KEAP1 labeling in vitro

KEAP1 was reconstituted in PBS containing 30% glycerol to a final concentration of 2 mg/mL. 1 μg KEAP1 in 50 μL PBS was incubated with the indicated concentrations of SA/succinyl coA at 37 °C for 1 hr, and then samples resolved by SDS-PAGE followed by Coomassie staining. The band containing KEAP1 was excised and submitted for MS/MS-based proteomics.

### Immunoblotting

Cells were collected by scraping in 250 μL ice cold 1x RIPA buffer with protease inhibitor. Lysates were sonicated using a tip sonicator (Branson) followed by clearing of insoluble material by centrifugation. The protein concentration of the supernatants was estimated by absorbance measurements on a Nanodrop instrument. 30 μg of lysate per lane was resolved using 4-12% Bis-Tris SDS PAGE gels (Invitrogen) and then transferred to PVDF membrane (Bio-Rad). Membranes were blocked in 5% non-fat dry milk (Biorad) in TBST (Tris-buffered saline with 0.1% Tween 20) for 1 hr at room temperature. Primary antibodies were incubated in milk in TBST overnight at 4 °C. Primary antibodies used here were anti-TUBG1 (Sigma, T6557, 1:2000), FLAG (Sigma, M2, 1:2000), anti-NRF2 (Proteintech, 66504, 1:2000), anti-succinyllysine (PTM Biolabs, PTM-401, 1:1000), anti-MYC (and anti-SUCLG2 (Bethyl Laboratories, A305-533A, 1:2000). After 5x TBST washes for 15-30 minutes, fluorophore-conjugated (1:2000, LI-COR) or HRP-conjugated (1:3000, ThermoFisher) secondary antibodies were incubated in milk for 1 hr and then washed again with TBST before visualizing on a LI-COR Odyssey fluorescent scanner or with film.

### Endogenous KEAP1 immunoprecipitation

HEK293T cells were plated, treated, and collected as described above. Lysate (2 mg/mL in 200 μL) was precleared with 20 μL of protein Pierce protein A/G magnetic beads (Thermo Fisher, 88802). 5 μg of anti-KEAP1 (Santa Cruz Biotechnologies, sc-365626) or anti-mouse IgG2b control (CST, 53484) was added to the lysate and incubated overnight at 4 °C. The following day, 20 μL of protein A/G beads were added and incubated with shaking for 3 hrs at 4 °C. After washing with lysis buffer, samples were boiled in 0.1% SDS and separated by SDS-PAGE.

### Statistical methods

Statistics were calculated in PRISM 9 (GraphPad, San Diego, CA). Data are presented as mean +/− SD and were analyzed by the student t-test or one- or two-way ANOVA, as indicated in the accompanying figure legends.

